# RUNX1 is regulated by androgen receptor to promote cancer stem markers and chemotherapy resistance in triple negative breast cancer

**DOI:** 10.1101/2022.11.12.516193

**Authors:** Natalia B Fernández, Sofía M Sosa, Justin T Roberts, María S Recouvreux, Luciana Rocha-Viegas, Jessica L Christenson, Nicole S Spoelstra, Facundo L Couto, Ana R Raimondi, Jennifer K Richer, Natalia Rubinstein

## Abstract

Triple negative breast cancer (TNBC) is an aggressive breast cancer subtype for which no effective targeted therapies are available. Growing evidence suggests that chemotherapy-resistant cancer cells with stem-like properties (CSC) may repopulate the tumor. Androgen receptor (AR) is expressed in up to 50% of TNBC, and AR inhibition decreases CSC and tumor initiation. Runt-related transcription factor 1 (RUNX1) correlates with poor prognosis in TNBC and is regulated by AR in prostate cancer. Our group has shown that RUNX1 promotes TNBC cell migration and regulates tumor gene expression. We hypothesized that RUNX1 is regulated by AR and that both may work together in TNBC CSC to promote disease recurrence following chemotherapy. Chromatin immunoprecipitation DNA-sequencing (ChIP-seq) experiments in MDA-MB-453 revealed AR binding to *RUNX1* regulatory regions. RUNX1 expression is upregulated by dihydrotestosterone (DHT) in MDA-MB-453 and in HCI-009 patient-derived xenograft (PDX) tumors (p<0.05). RUNX1 is increased in a CSC-like experimental model in MDA-MB-453 and SUM-159PT cells (p<0.05). Inhibition of RUNX1 transcriptional activity reduced the expression of CSC markers. Interestingly, RUNX1 inhibition reduced cell viability and enhanced paclitaxel and enzalutamide sensitivity. Targeting RUNX1 may be an attractive strategy to potentiate the anti-tumor effects of AR inhibition, specifically in the slow growing CSC-like populations that resist chemotherapy leading to metastatic disease.

## INTRODUCTION

Triple negative breast cancer (TNBC) is a heterogeneous disease that includes all breast cancer subtypes that have no expression of estrogen and progesterone receptors and no amplification of human epidermal growth factor receptor 2 (HER2) [1]. Accounting for 15–20% of all breast cancers, TNBC is more prevalent in younger women and women of African and Hispanic descents, as reviewed in Zagami et al. (2022) [2]. TNBC has been recently subdivided into four subtypes on the basis of gene expression profiles: basal-like immunosuppressed (BLIS), immunomodulatory (IM), luminal androgen receptor (LAR) and mesenchymal (MES) [3]. Despite major advances in breast cancer treatments, the heterogeneity of TNBC is still a challenge since there are few targeted therapy options, leaving surgery, chemotherapy, and radiation as the first line of treatment [4]. Unfortunately, 35% of the tumors will escape these strategies leading to a high rate of rapid recurrence as metastatic disease [5]. The lack of better precision therapeutic targets is an unmet need for these tumors and contributes to a more aggressive behavior for this tumor subtype [4,6,7]. Studies suggest that chemotherapy-resistant cancer cells with stem-like properties may repopulate the tumor locally or cause recurrence as metastatic disease [4,8–10]. Thus, therapies that target the cancer stem cell (CSC)-like population in combination with chemotherapy to prevent rapidly dividing cells may impair tumor recurrence. Growing evidence suggests that chemotherapy resistance is associated with a more slowly dividing CSC subpopulation [4,8–10]. The LAR subtype represents 20-40% of TNBCs and is characterized by expression of the androgen receptor (AR), a luminal-like pattern of gene expression [3] and reduced pathologic complete response to neoadjuvant chemotherapy [11]. AR has emerged as a potential therapeutic target in breast cancer and its efficacy in AR^+^-TNBC (Stage I-III) patients is currently under evaluation, based on a protocol combining enzalutamide (Enza) and paclitaxel (Px) before surgery (clinical trial NCT02689427). Indeed, AR inhibition significantly reduces baseline proliferation, anchorage-independent growth, migration and invasion, and increases apoptosis in AR^+^-TNBC lines [12]. *In vivo*, Enza significantly decreases viability of SUM-159PT and HCC1806 xenografts [12]. Moreover, AR supports CSC-like properties, including anchorage-independent survival and mammosphere formation [13]. Pretreatment with Enza reduces tumor volume and viability when administered simultaneously or subsequently with Px and simultaneous treatment suppressed tumor recurrence more effectively in mice after drug cessation [13]. Moving forward, a biomarker for the selection of patients with TNBC suitable for treatment with AR inhibitors is a major unmet need [7]. Nevertheless, a precise understanding of the mechanism of androgen action in this disease remains a challenging puzzle.

The runt-related transcription factor (RUNX) family of transcription factors regulate a plethora of developmental processes including cell growth, differentiation and lineage specification [14–16]. During mammary development, RUNX factors are important for the maintenance of mammary epithelium homeostasis [17,18]. In human breast cancer, RUNX1 activity is still a matter of debate, and little is known about its role in tumor progression. Accumulating evidence strongly suggests that RUNX1 promotes tumor aggressiveness in TNBC, while functioning as tumor suppressor in ER^+^ breast cancer [17,19–26].

According to Cancer Genome Atlas Network (2012) and Caldas analysis [27], the *RUNX1* gene is mutated only in Luminal A/B tumors. Moreover, in the Catalogue of Somatic Mutations in Cancer, *RUNX1* is included in the top 20 mutated genes. However, no mutation has been reported in TNBC samples suggesting that there might be a selective pressure to maintain wild-type protein in TNBC (COSMIC, http://cancer.sanger.ac.uk/cosmic/). Additionally, RUNX1 has an independent prognostic indicator of poor patient outcomes in TNBC [20]. RUNX transcription factors and their coregulator, core binding factor beta (CBFβ), promote phenotypic plasticity and are essential for maintaining the mesenchymal and invasive phenotype [23]. Our group showed that RUNX1 regulates R-Spondin 3 (RSPO3), promoting tumor growth and motility [21,28]. Strikingly, it has been reported that RUNX1 is involved in the differentiation or reduction of normal and tumoral ER^+^ mammary stem cells [24,29,30] and in the proliferation of mesenchymal prostate stem cell proliferation [31]. However, there is no data on its role in TNBC-CSC yet. It has been reported that AR binds to the *RUNX1* promoter in prostate cancer cell lines, but the relationship between these two proteins in breast cancer is still uncertain [32].

Our goal was to investigate the relevance of RUNX1 in AR^+^-TNBC tumors. Our hypothesis is that RUNX1 is regulated by AR activation to promote CSC enrichment, leading to a chemoresistant population capable of surviving and metastasizing to distant organs. Activation of AR by dihydrotestosterone (DHT) induces *RUNX1* gene expression *in vitro* and *in vivo* in AR^+^-TNBC PDX tumor samples. By inhibiting RUNX1 transcriptional activity we determined that it is required to induce CSC genes and to enhance chemoresistance. Our results show, for the first time, that AR induces RUNX1 expression in TNBC cell lines promoting a CSC phenotype enrichment and increased chemoresistance. Furthermore, these data suggest that AR and RUNX1 might work together to promote tumor progression and be useful for clinical therapeutic decision-making in AR^+^-TNBC.

## MATERIAL AND METHODS

### Cell lines and reagents

MDA-MB-453 cells were purchased from the ATCC and maintained in DMEM high glucose medium with 10% fetal bovine serum (FBS, Sigma). SUM-159PT were obtained from the University of Colorado Cancer Center (UCCC) Tissue Culture Core (Aurora, CO) and maintained in Ham’s F-12 with 5% FBS, 1% HEPES, 1 μg/mL hydrocortisone and 5 μg/mL insulin. BT-549 cells, purchased from the ATCC, were grown in RPMI 1640 medium with 10% FBS, nonessential amino acids (NEAA) and 5 μg/mL insulin supplementation. MDA-MB-231 cells were grown in RPMI 1640 medium with 10% FBS. All cells were grown in the presence of 1% streptomycin and amphotericin B (Gibco) and maintained at 37°C in a humidified incubator containing 95% air and 5% CO_2_. Only cells under ten passages were used in this study. All cell lines were provided by the laboratory of Dr. Jennifer Richer, see Barton 2017 for details [13].

Dihydrotestosterone (DHT, Sigma-Aldrich) was diluted in ethanol, Enzalutamide (Enza, Sigma-Aldrich #PHB00235) in DMSO, Paclitaxel (Px, Cell Signaling #98075) in DMSO and RUNX1 inhibitors AI-10-104 (Glixx Laboratories GLXC-20705) and AI-10-49 (Glixx Laboratories GLXC-07203) in DMSO. All experiments that included DHT and/or Enza treatment were conducted in charcoal-stripped serum.

### Forced suspension culture

Poly-2-hydroxyethyl methacrylate (poly-HEMA, Sigma) was prepared at a concentration of 12 mg/ml in 95% ethanol. Culture plates were incubated with poly-HEMA overnight to allow ethanol evaporation. Plates were washed with PBS prior to use.

### Quantitative RT-PCR

RNA was isolated by TRI reagent (MRC) and cDNA was synthesized from 1 μg total RNA, using M-MLV reverse transcriptase enzyme (Promega). SYBR Green quantitative gene expression analysis was performed using Taq polymerase (Thermo Fisher) in a StepOne instrument (Applied Biosystem). Relative gene expression was calculated using the 2 –ΔΔCt method and values were normalized to GAPDH. Primer sequences are listed in supplementary Table S4a.

### Western blot

Protein extracts were prepared in a cell lysis buffer and denatured at 95 °C for 10 minutes, separated on SDS-PAGE gels and transferred to nitrocellulose membranes (Bio-rad). After blocking in 5% non-fat milk in T-TBS, membranes were incubated overnight at 4 °C with primary antibodies in 0.5% BSA or 2% non-fat milk in T-TBS: anti-AR (1:1000 dilution; Santa Cruz Biotechnology #7305), anti-RUNX1 (1:1000 dilution; Cell Signaling Technology #4334), anti-SOX4 (1:1000 dilution; Abcam #52043), anti-Tubulin (1:10000, Sigma #T5168) and anti-GAPDH (1:5000; Santa Cruz Biotechnology #32233). Secondary antibody incubation was performed at room temperature for 1 hour: anti-mouse (1:5000, Li-Cor #926-32213 or Li-Cor #926-68070) or anti-rabbit (1:5000, Li-Cor #926-32210). Membranes were then scanned using Odyssey Imaging System and analyzed with Image Studio Ver 5.2 software (Li-Cor).

### Chromatin immunoprecipitation

For AR ChIP-seq, MDA-MB-453 cells were grown in charcoal-stripped serum media for a total of 72 hrs before treatment. Twenty-four hours prior to treatment, cells were trypsinized and equal cell numbers were plated on control tissue culture dishes (attached) or poly-Hema coated dishes (suspended). Cells were treated with DMSO (vehicle control), DHT (10 nM), or DHT+Enza (10 μM) for 4 hours, followed by fixation in 1% formaldehyde. ChIP-seq was performed as previously reported [43]. Chromatin was sonicated using an Epishear Probe Sonicator (Active Motif) for 4 minutes (cycles of 30 seconds with 30 seconds of rest in between) at 40% power. AR antibody H-280 (Santa Cruz) was utilized for immunoprecipitation and libraries were sequenced on an Illumina HiSeq 2500.

RUNX1 ChIP in MDA-MB-231 cells was performed as previously described [21] using anti-RUNX1 (Abcam #23980) and anti-IgG (Abcam #46540, negative control). Primer sequences are available in supplementary Table S4b.

### ALDEFLUOR

The ALDEFLUOR assay (Stem Cell Technologies) was performed per the manufacturer’s protocol and as previously reported [13]. ALDEFLUOR-positive and - negative cell populations were sorted with the assistance of the University of Colorado Cancer Center Flow Cytometry Shared Resource on the MoFlo XDP100 cell sorter (Beck- man Coulter Life Sciences).

### Crystal Violet

Cells were plated in 96-well plates in quadruplicate or quintuplicate and treated with increasing concentrations of AI-10-104/-49 (for dose-response curves) or with Px, Enza, AI-10-104/-49 alone or in combination (for cytotoxicity assays). After 3 days of drug treatments, cells were fixed with 10% formalin, stained with 0.1% crystal violet, and then solubilized with 10% acetic acid. Absorbance was measured at 540 nm. In parallel, another plate with an increasing number of cells was prepared in order to generate a calibration curve. Data are presented as percentage of cell viability relative to control treated cells (vehicle, DMSO).

### MTT

Cells were plated in 96-well plates in quintuplicate and treated with Px, Enza, AI-10-104/-49 alone or in combination. After 3 days of drug treatments, 0.5 μg/μl thiazolyl blue tetrazolium bromide (Sigma #M5655) was added at each well and incubated for 4 hours at 37 °C. Then, 0.01N Iso-propanol was used to dissolve the formazan crystals and absorbance at 570 nm was measured. Data are presented as the relative absorbance to control treated cells (vehicle, DMSO).

### Statistical analysis

Statistical significance was evaluated using two-tailed unpaired Student t-tests or one-way ANOVA followed by Tukey contrast with GraphPad Prism 9 software. All experiments were performed at least 3 times before analyzing the statistical significance. A *p* value of less than 0.05 was considered statistically significant.

## RESULTS

### Androgen receptor (AR) regulates *RUNX1* gene expression

To evaluate the capacity of AR to regulate *RUNX1* gene expression, an AR ChIP-seq assay was performed in MDA-MB-453, a representative LAR TNBC cell line according to gene expression profile [33–35]. AR binds to the *RUNX1* promoter and four intronic loci within the *RUNX1* gene body **(Figure 1A)**. Treatment for 24 hours with DHT (an AR agonist) increases AR recruitment to *RUNX1* in all the sites and this effect was blocked in the presence of the AR antagonist Enza. Moreover, DHT treatment upregulates *RUNX1* mRNA **(Figure 1B)** and protein **(Figure 1C)** levels, an effect that is also blocked when Enza is added. Future analyses are necessary to determine the relative functional contributions of these regions.

**FIGURE 1.**
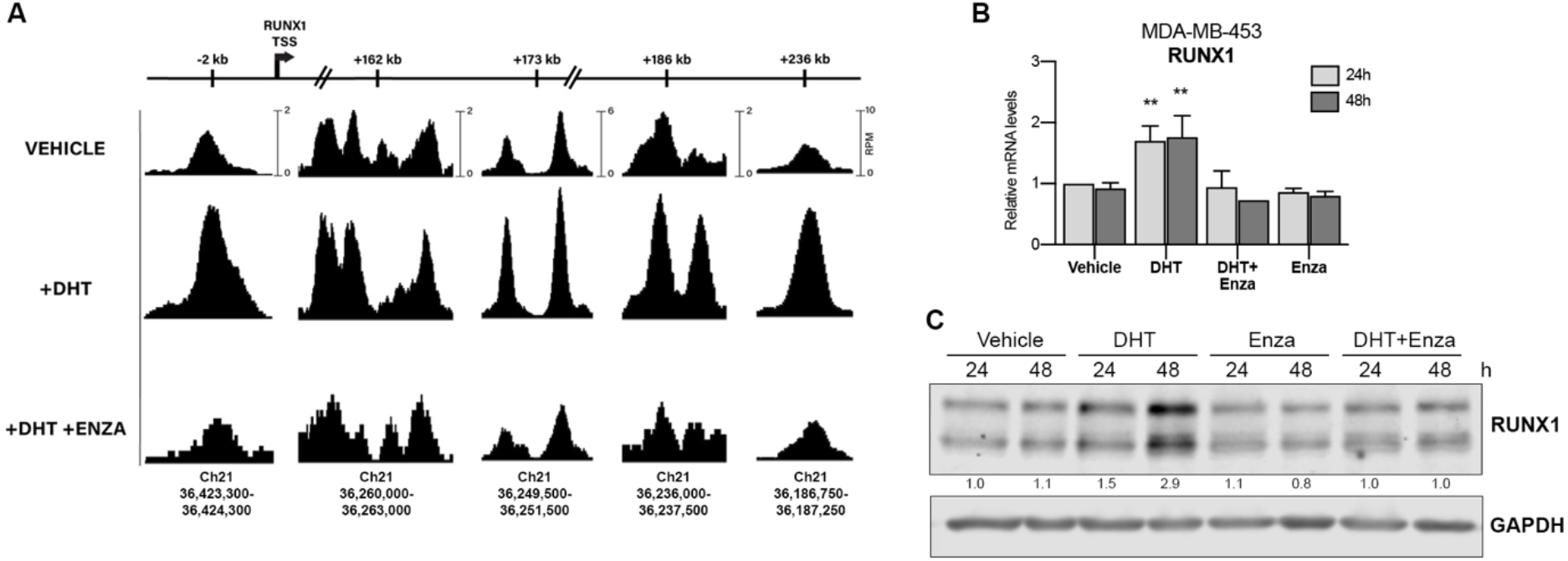
AR regulates RUNX1 expression. **A)** MDA-MB-453 cells were treated with either vehicle (DMSO+ethanol), DHT (10 nM) or DHT+Enza (20 uM) for 24 hrs. AR ChIP-seq was performed and the RUNX1 promoter and intronic regions were analyzed. The scale bars at the top are labeled for each locus to indicate the relative distance from the canonical RUNX1 transcription start site (TSS). The chromosomal coordinates of each peak are shown below. RUNX1 qPCR **(B)** and Western blot **(C)** were performed in MDA-MB-453 cells treated with either vehicle (DMSO+ethanol), DHT (10 nM), Enza (20 uM) or both for 24 and 48 hrs. GAPDH was used as a housekeeping control. ** p<0.01.

This modulation is also observed in RNA-seq of HCI-009, an AR^+^ PDX tumor grown in mice with or without DHT, which showed significant upregulation of *RUNX1* **(Table 1a)**. Furthermore, scRNA-seq analysis between AR^High^ versus AR^Low^ MDA-MB-453 cell populations showed significantly higher *RUNX1* expression in the AR^High^ cells [36] **(Table 1b)**. All together these data show that AR positively regulates RUNX1 in AR^+^-TNBC.

**Table 1a.** Gene expression changes following DHT treatment in HCI-009 PDX.

| Gene | Common Name | Fold change | p value | Adjusted p value |
| --- | --- | --- | --- | --- |
| ENSG00000159216 | *RUNX1* | 1,84 | 1,27E-03 | 2,27E-02 |

**Table 1b.**
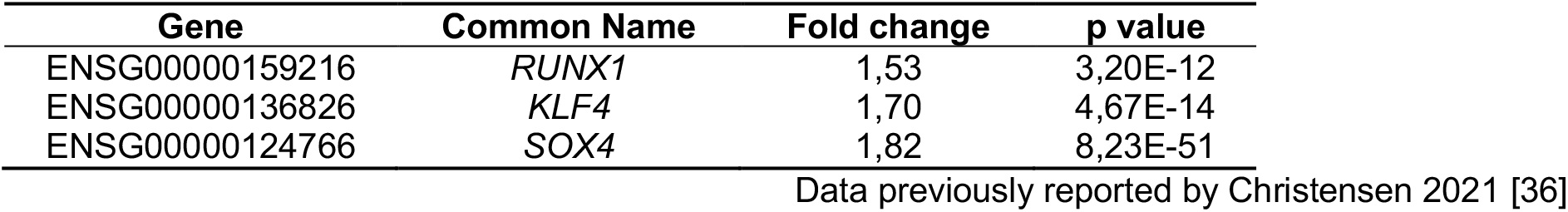
Gene expression differences between AR^High^ versus AR^Low^ MDA-MB-453 cells.

### RUNX1 is upregulated in a circulating tumor cell model and contributes to a CSC phenotype

Circulating tumor cells (CTC) with CSC-like phenotype have been described as the major source of clinical tumor recurrence [37,38]. To evaluate the contribution of RUNX1 in the physiology of this subpopulation we used a forced suspension *in vitro* cell culture model since AR^+^-TNBC cells grown in these conditions’ express higher levels of AR and CSC markers such as CD22/CD44 and increased ALDH activity levels than their attached counterparts [13]. We observed that the AR binding sites in the *RUNX1* gene were conserved when MDA-MB-453 cells were cultured in forced suspended conditions and responded to DHT and Enza treatment **(Figure S1A)**, like in attached conditions. We found that *RUNX1* mRNA is upregulated in MDA-MB-453 cells surviving in forced suspension culture over time **(Figure 2A)** and that treatment with DHT and Enza also modulates RUNX1 protein levels in these conditions **(Figure 2B** and **Figure S1B)**. The rise of RUNX1 expression in this CTC model was accompanied by an increase of CSC markers, such as krüppel-like factor 4 (*KLF4*) and octamer-binding transcription factor 4 (*OCT4*) **(Figure 2C)**.

**FIGURE 2.**
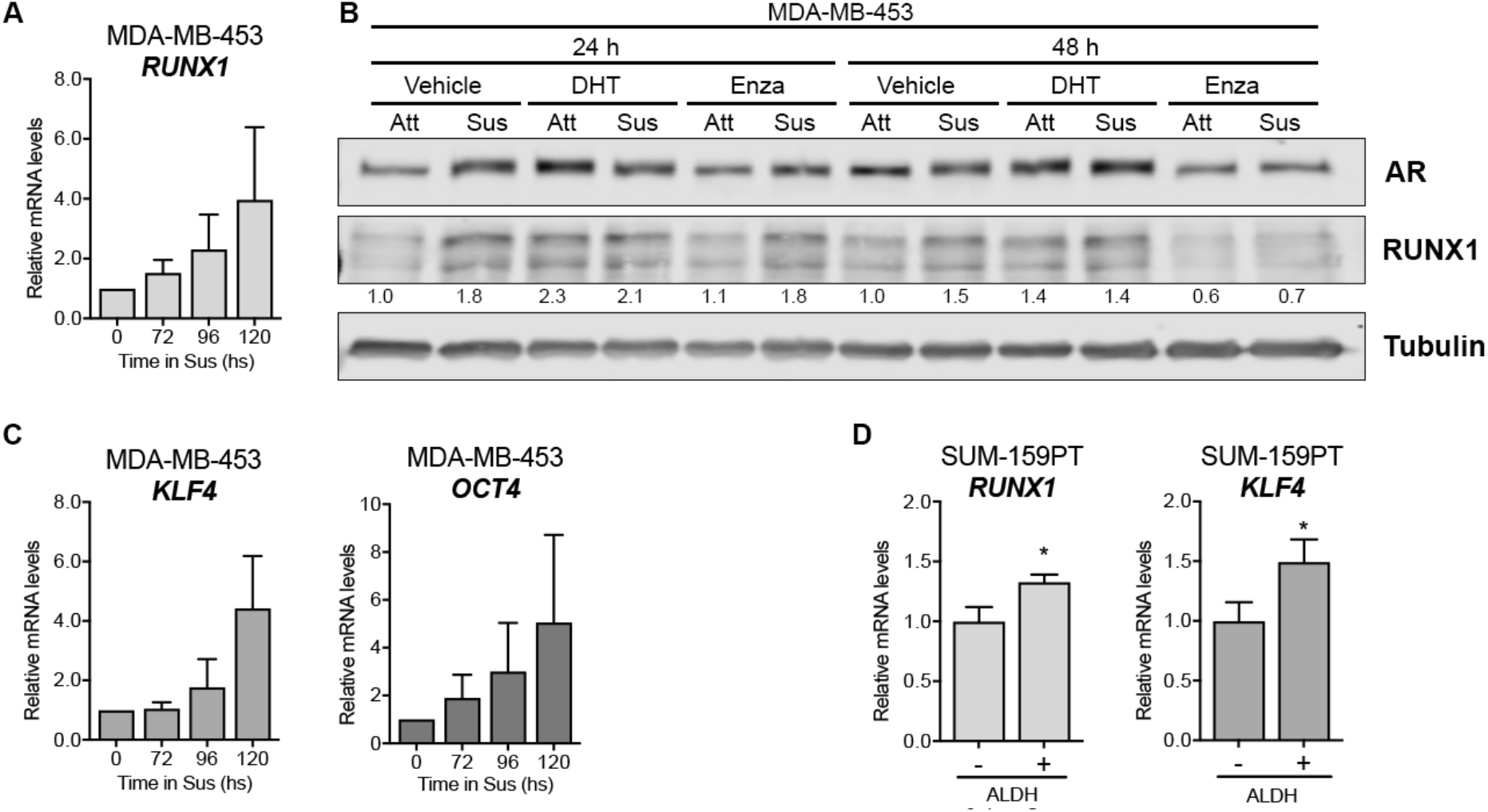
RUNX1 is upregulated in CSC-like subpopulations. mRNA levels of *RUNX1* **(A)**, *KLF4* and *OCT4* **(C)** after 72, 96 and 120 hrs in forced suspension (Sus) were determined by qPCR. *GAPDH* was used as a housekeeping control. **B)** AR and RUNX1 Western blot of MDA-MB-453 cells culture in attached (Att) or forced-suspension conditions (Sus) and treated with either vehicle (DMSO+ethanol), DHT (10 nM) or Enza (20 uM) for 24 and 48 hrs. Tubulin was used as a housekeeping control. **D)** SUM-159PT were cultured in forced-suspension conditions for 3 days and then were sorted using ALDEFLUOR assay. mRNA was prepared from ALDH^-^ and ALDH^+^ subpopulations and *RUNX1* and *KLF4* levels were evaluated by qPCR. *GAPDH* was used as a housekeeping control. * p<0.05.

It was previously reported that SUM-159PT cells cultured for 3 days in forced suspension increase the population of aldehyde dehydrogenase-positive (ALDH^+^) cells by 60% and express significantly higher levels of AR [13]. Since ALDH is a CSC marker we evaluated RUNX1 expression in this ALDH^+^ subpopulation. ALDH^+^ cells express significantly higher levels of *RUNX1* and *KLF4* after 3 days in forced suspension compared to levels in ALDH^-^ cells, which is consistent with a CSC-like phenotype (**Figure 2D**). *AR* expression was examined as an internal positive control **(Figure S1C)**.

To investigate the contribution of RUNX1 in modulating the CSC phenotype in this CTC model we inhibited its transcriptional activity and measured the expression of CSC gene markers. RUNX1 commercial inhibitors AI-10-104 and -49 both downregulated *KLF4* and *OCT4* gene expression in MDA-MB-453 cells cultured in forced suspension **(Figure 3)**.

**FIGURE 3.**
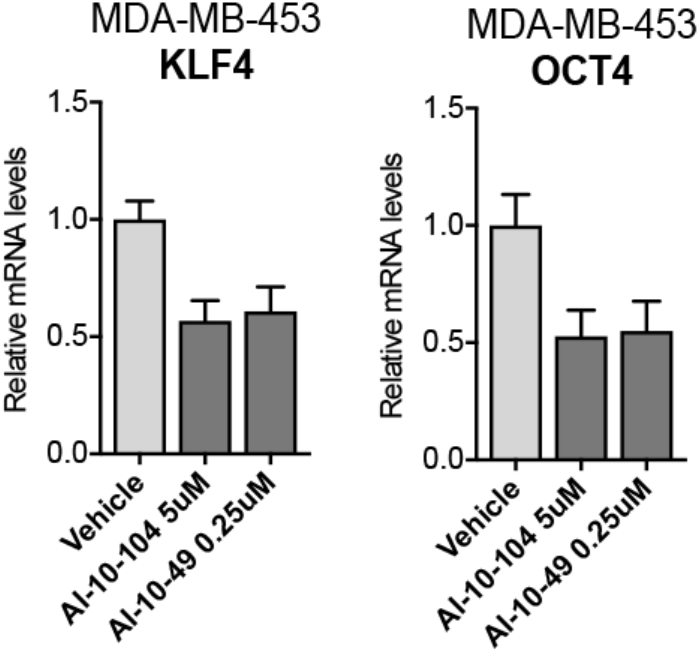
RUNX1 is required for the generation of CSC-like phenotype. MDA-MB-453 cells were cultured in forced-suspension for 4 days and treated for one extra day with AI-10-104 (5 uM) or AI-10-49 (0.25 uM). mRNA levels of *KLF4* and *OCT4* were determined by qPCR. *GAPDH* was used as a housekeeping control.

In order to further investigate RUNX1 involvement in the CSC phenotype we examined other relevant putative target genes. A previous RUNX1 ChIP assay performed by our group in MDA-MB-231 cells revealed that RUNX1 binds to different tumor related genes [21]. Here we report that RUNX1 also binds to the *SOX4* gene promoter **(Figure 4A)**. SOX4 is an interesting transcription factor because it has been implicated in breast cancer Epithelial-Mesenchymal Transition (EMT) [39], metastasis [40] and drug resistance in other tumors such as colon cancer [41]. To evaluate the ability of RUNX1 to regulate SOX4 expression, MDA-MB-453 and SUM-159PT cell lines were treated with AI-10-104 for 24 hrs. SOX4 protein is decreased in these cell lines by RUNX1 inhibition (**Figure 4B**). Moreover, when these cell lines were cultured in forced suspension for 4 days and treated with the RUNX1 inhibitors for 24 hours, SOX4 protein was also downregulated **(Figure 4C)**.

**FIGURE 4.**
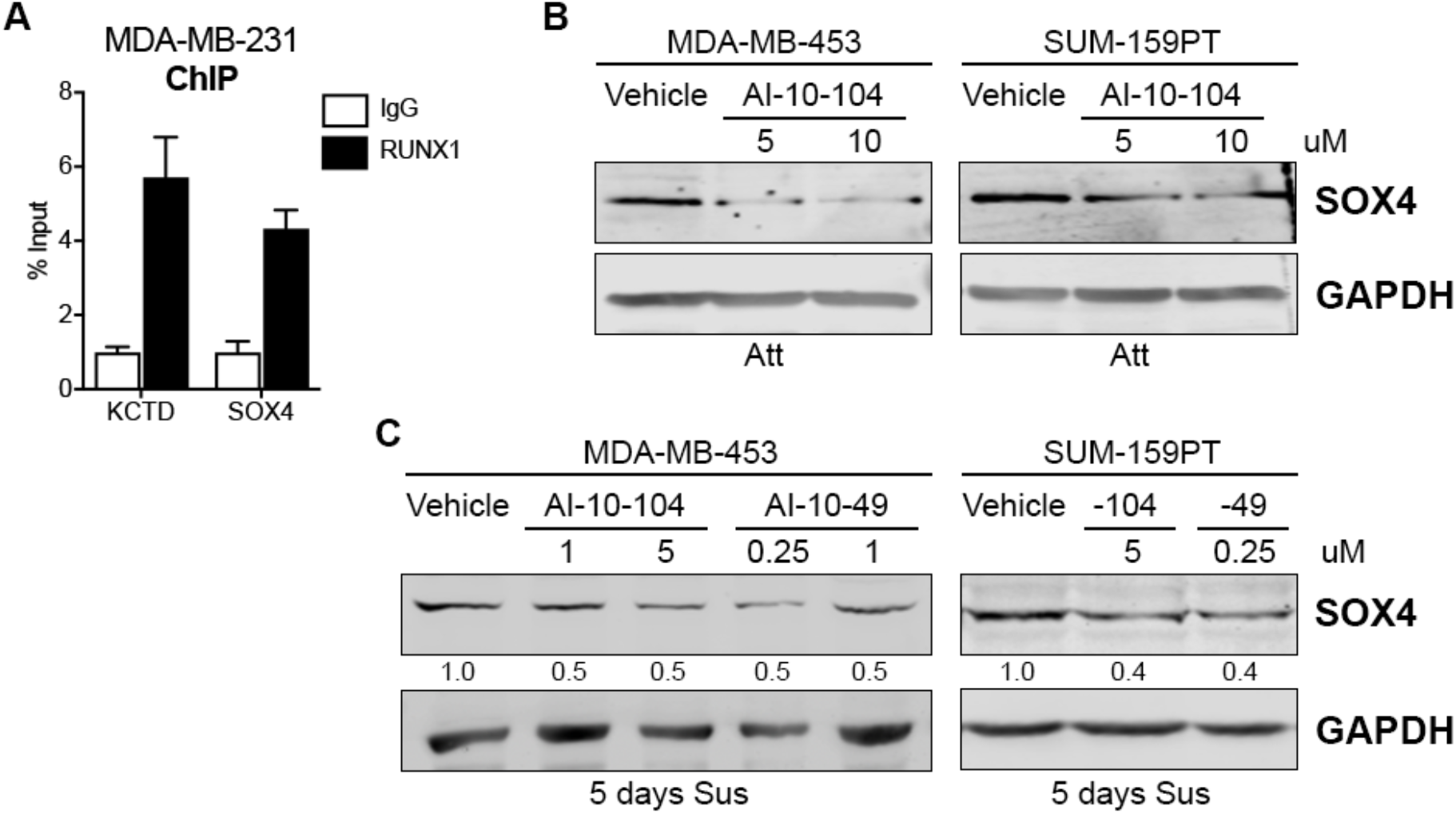
RUNX1 binds to SOX4 promoter and regulates its expression. **A)** RUNX1 ChIP assay was performed in MDA-MB-231 and *SOX4* promoter primers were designed to determine RUNX1 binding by qPCR. KCTD was used as a positive control. **B)** SOX4 Western blot was performed in MDA-MB-453 and SUM-159PT cultured in attached (Att) conditions and treated with AI-10-104 (5 or 10 uM) for 24 hrs. *GADPH* was used as a housekeeping control. **C)** MDA-MB-453 and SUM-159PT were cultured in forced-suspension (Sus) for 4 days and treated for one extra day with AI-10-104 (1 or 5 uM) or AI-10-49 (0.25 or 1 uM). *GADPH* was used as a housekeeping control.

Interestingly, *KLF4* and *SOX4* mRNAs were also upregulated in AR^High^ cells compared to AR^Low^ cells (**Table 1b**, [36]), suggesting that, even under standard adherent culture conditions, these genes might be involved in the AR/RUNX1 axis that drives tumor cell fate.

### Loss of RUNX1 transcriptional activity reduces AR^+^-TNBC viability and enhances drug sensitivity

Since the CSC and EMT phenotype are involved in drug resistance in TNBC [4,8–10], we explored RUNX1 involvement in chemoresistance. To investigate the role of RUNX1 in response to chemotherapeutic drugs, AR^+^-TNBC cell lines were treated with the RUNX1 transcriptional activity inhibitors AI-10-104 and AI-10-49, the AR antagonist Enza, and the chemotherapeutic drug Px. It is known that inhibition of AR combined with chemotherapy resulted in a more effective outcome than chemotherapy alone *in vitro* and *in vivo* in preclinical mouse models [12]. This combination is currently being tested in a clinical trial (NCT02689427) based on these preclinical data. Reduction in RUNX1 transcriptional activity decreases MDA-MB-453 and SUM-159PT cell viability in a dose dependent manner (**Figure 5A** and **5B**). Inhibition of RUNX1 also reduces tumor cell colony formation (**Suppl Figure S2A**). Importantly, reduction of RUNX1 transcriptional activity with either AI-10-104 or AI-10-49 significantly increased sensitivity to Px and Enza in standard tissue culture conditions after treatment for 72 hrs **(Figure 5C-F** crystal violet assays and **S2B** MTT assay). To further explore the role of RUNX1 in CSC/CTC drug resistance, anchorage-independent cells were treated with Enza, Px and RUNX1 inhibitors for cell survival evaluation by MTT assay. A reduction in RUNX1 transcriptional activity significantly improves the efficacy of Px and Enza in forced suspension culture (**Figure 6**). Under these conditions we had to use higher doses of all the drugs, including RUNX1 inhibitor, to achieve a reduction in cell viability rate similar to the one obtained in standard adherent cultures (**Figure 6**), validating the already known concept that CSC populations are more resistant to treatment [42].

**FIGURE 5.**
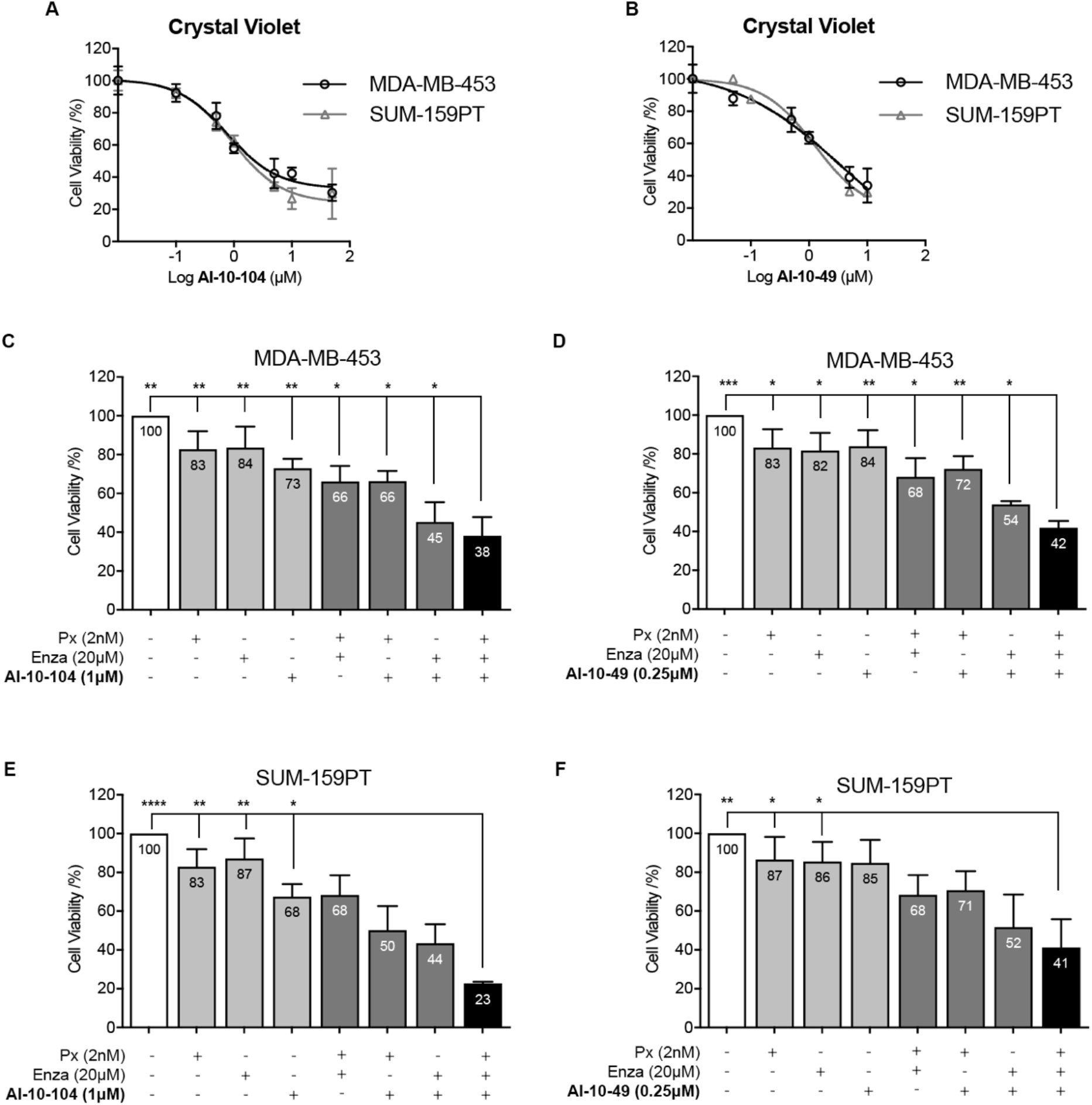
Inhibition of RUNX1 transcriptional activity reduces cell viability and enhances drugs’ cytotoxic effects. **A-B)** MDA-MB-453 and SUM-159PT cells were cultured for 72 hrs and treated with increasing concentration of AI-10-104 **(A)** or AI-10-49 **(B)**. MDA-MB-453 **(C, D)** and SUM-159PT **(E, F)** were treated with 2 nM Px, 20 μM Enza and 1 μM AI-10-104 **(C, E)** or 0.25 μM AI-10-49 **(D, F)** or all possible combinations for 72 hrs. In all cases, cell viability was determined by crystal violet assay (absorbance at 570 nm) using a calibration curve. Percentage of cell viability was calculated and expressed as relative to vehicle treatment (DMSO, 100%). Statistical differences are shown only for the combination of the 3 drugs vs the rest of the treatments, for more details see Suppl Table S1 and S2. * p<0.05, ** p<0.01, *** p<0.001 and **** p<0.0001.

**FIGURE 6.**
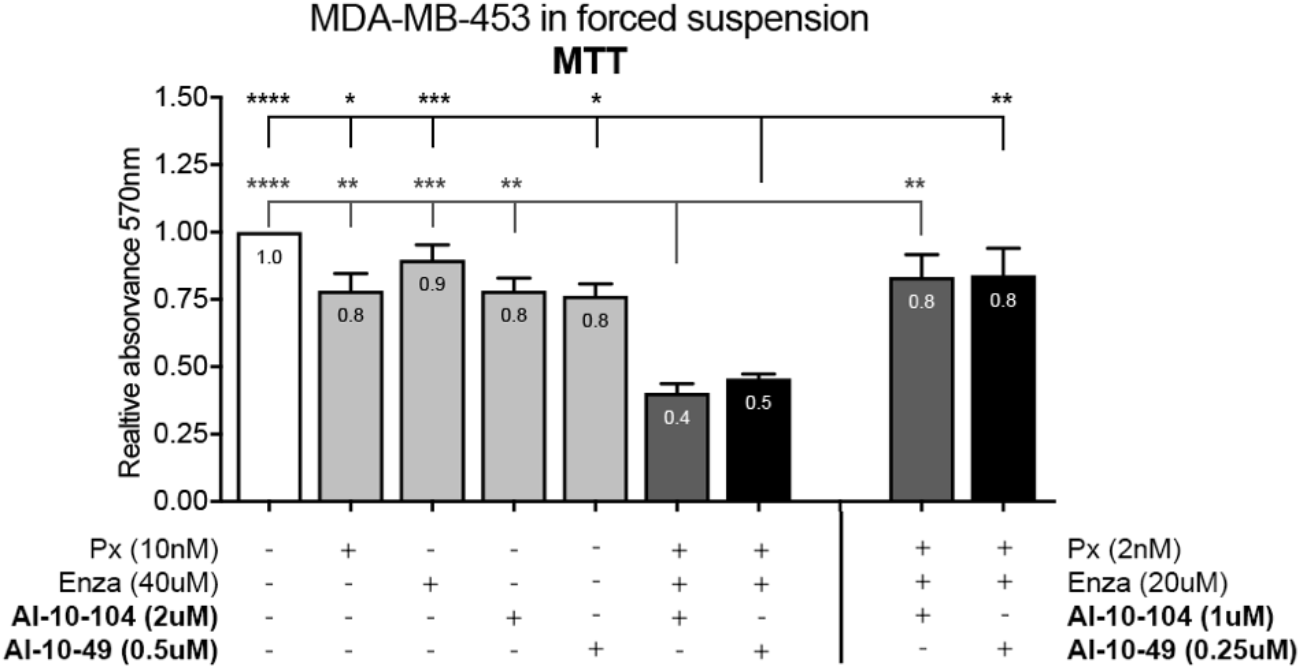
Reduction of RUNX1 transcriptional activity enhances enzalutamide and paclitaxel treatment in CSC-like cells. MDA-MB-453 cells were cultured for 72 hrs in forced-suspension and treated with either 10 nM Px, 40 μM Enza and 2 μM AI-10-104 / 0.5 μM AI-10-49 (right panel) or the doses used for attached conditions 2 nM Px, 20 μM Enza and 1 μM AI-10-104 / 0.25 μM AI-10-49 (left panel) for 72 hrs. Cell viability was determined by MTT and the results are expressed as the relative absorbance (570 nm) to control treatment (DMSO, 1.00). Statistical differences are shown for the combination of the 3 drugs vs the rest of the treatments (AI-10-104 in dark gray and AI-10-49 in black), for more details see Suppl Table S3. * p<0.05, ** p<0.01, *** p<0.001 and **** p<0.0001.

Taken together, these results strongly suggest that RUNX1 transcriptional activity is necessary for TNBC cells to survive chemotherapy. More importantly, they show that reducing RUNX1 transcriptional activity may be an opportunity to improve the sensitivity of CSC/CTC to chemotherapy leading to a potential reduction in metastasis or tumor recurrence in AR^+^-TNBC.

## DISCUSSION

Here we show for the first time that AR is a direct positive regulator of *RUNX1* gene expression in AR^+^-TNBC and that RUNX1 transcriptional activity is involved in the CSC-like phenotype and chemoresistance, contributing to tumor progression in this aggressive breast cancer subtype.

One of the current principal clinical challenges in TNBC is the presence of inter- and intratumoral heterogeneity that hinders decision making and drives the lack of therapeutic efficacy [44,45]. Therefore, identification of better markers could reveal more accurate therapeutic strategies. Although several findings support a role for the androgen/AR axis in breast cancer, its involvement in the pathogenesis and progression of this cancer remains debated. The predictive and prognostic role of AR in TNBC is still clinically undetermined [46]. Accumulating data suggests that even if a protein is indicative of a more well-differentiated tumor, it can serve as a therapeutic target if the tumors are dependent upon it for growth (as is well acknowledged for targeting ER). It is clear that AR^+^-TNBC has a poor pathological complete response (pCR) [11], and anti-androgen therapy (to kill the slow growing cells) combined with standard chemotherapy drugs, such as paclitaxel (to kill rapidly dividing tumor cells), showed promising results in breast cancer preclinical models [12,13]. Interestingly, a clinical trial is currently underway in AR^+^-TNBC patients using this drug combination (NCT02689427). The determination of better biomarkers for the selection of patients suitable for treatment with AR inhibitors is a major unmet need.

We determined that RUNX1 may be an appropriate marker for anti-androgen treatment in AR^+^-TNBC and also a potential therapeutic target by itself. Blocking RUNX1 transcriptional activity has already been tested in a preclinical model of leukemia showing promising results [47]. In Figure 3 we show that RUNX1 transcriptional activity inhibition has a strong negative effect on *KLF4* and *OCT4* gene expression. Both genes have been described as determinant factors involved in CSC development [48] and in TNBC CSC enrichment [49,50]. Since four RUNX1 binding sites were identified in the *KLF4* promoter in a human leukemia cell model [51] and KLF4 is an AR gene target in breast cancer cell lines [52], more experiments are needed to define the principal source of *KLF4* gene expression activation.

Furthermore, in Figure 4 we report that RUNX1 binds to the *SOX4* gene promoter and regulates its expression in standard and suspended conditions, suggesting that SOX4 could be one of the RUNX1 mediators in promoting CSC and/or chemoresistance in our model. In line with this, it has been described that stable overexpression of SOX4 in immortalized, non-transformed RWPE-1 prostate cells enables anchorage independent growth and colony formation in soft agar [53]. It has been recently demonstrated that combined inhibition of Wnt signaling and SOX4 inhibits proliferation and migration and induces apoptosis of TNBC BT-549 cells [54]. In supplementary Figure S2C we show that inhibition of RUNX1 also potentiates Px and Enza toxicity in this AR^+^-TNBC cell line. Several studies have indicated that SOX4 also plays a critical role in EMT regulation, which can facilitate metastasis and chemoresistance in carcinomas [55]. Indeed, other groups have shown that SOX4 overexpression induces EMT in breast cancer cells [39], which in turn upregulates stem cell markers and enhances mammosphere formation [56]. In addition, SOX4 involvement in drug resistance has been described in colon and cervical cancer [41,57]. The present observation that RUNX1 regulates SOX4 unravels a potential mechanism by which RUNX1 regulates EMT in TNBC cell lines, previously reported in Ran (2020) [23]. In sum, our data strongly suggests that RUNX1 may be required for the enrichment of CSC in AR^+^-TNBC cell lines.

In contrast to TNBC, RUNX1 and CBFβ have been described as tumor suppressor genes involved in reduced tumor growth and impaired EMT and CSC generation in ER^+^ breast cancer [26,58]. Remarkably, cumulative evidence supports the concept that the role of RUNX1 in breast cancer depends on hormone receptor context [17,20– 23].

Collectively, our results reveal that reduction of RUNX1 transcriptional activity significantly increases sensitivity to chemotherapy in AR^+^-TNBC cell lines, in both standard culture conditions or in forced suspension. Pharmacologic inhibition of RUNX1 significantly enhances the previously described combination treatments, such as Px and Enza. Remarkably, we observed that cells grown in suspended conditions (CSC-like phenotype) need higher concentrations of drugs than the attached ones to generate a similar toxic effect. This observation is supported by accumulated evidence suggesting that CSC are the remaining population that survives drug treatments and regenerates tumors [48]. RUNX1 participation in chemotherapy drug response has also been reported in ovarian cancer [59], glioblastoma multiforme [60] and colorectal cancer [61], suggesting that this function could be a general molecular mechanism of action that favors tumor aggressiveness against cytostatic drug treatment. *In vivo* experiments have yet to be performed to continue exploring this concept in triple negative breast cancer.

Finally, it has been described that the tumor microenvironment is relevant in TNBC tumor progression and chemoresistance [62]. Recently, Halperin (2022) reported that RUNX1 expression is upregulated in cancer-associated fibroblasts and that the RUNX1 signature is associated with poor breast cancer outcome [63]. This recent evidence suggests that blocking the transcriptional activity of RUNX1 *in vivo* could simultaneously reduce tumor growth and increase its sensitivity to drugs, as well as weaken the pro-tumorigenic effect of its microenvironment.

## AUTHOR CONTRIBUTION

Conceptualization, NR and NBF; methodology, JR, NBF, SMS, JTR, LRV, MSR, FLC, JLC, NSS, ARR and NR; formal analysis, NBF, SMS, LRV, JTR and NR; writing— original draft preparation, NBF and NR; writing—review and editing, NR, NBF and JKR; visualization, NBF and SS.; supervision, NR.; project administration, NR.; funding acquisition, NBF, JKR and NR. All authors have read and agreed to the published version of the manuscript.

## ACKNOWLEDGMENTS

The authors would like to thank Pilar Bastida for her support and MACMA Foundation for advocate point of view support.

## SUPPLEMENTAL FIGURES AND TABLE LEGENDS

**Supplementary Figure S1. A)** MDA-MB-453 cells were cultured in forced suspended conditions and treated with either vehicle (DMSO+ethanol), DHT (10 nM) or DHT+Enza (20 uM) for 24 hrs. AR ChIP-seq was performed and the RUNX1 promoter and intronic regions were analyzed. The scale bars at the top are labeled for each locus to indicate the relative distance from the canonical RUNX1 transcription start site (TSS). **B)** AR and RUNX1 Western blot of MDA-MB-453 cells in attached (Att) or forced-suspended conditions (Sus) and treated with either vehicle (DMSO+ethanol), DHT (10 nM) or Enza (20 uM) for 72 hrs. Tubulin was used as a housekeeping control. **C)** SUM-159PT were cultured in forced-suspended conditions (Sus) for 3 days and then were sorted using the ALDEFLUOR assay. mRNA was prepared from ALDH^-^ and ALDH^+^ subpopulations and AR levels were evaluated by qPCR. ** p<0.01.

**Supplementary Figure S2. A)** SUM-159PT were cultured in soft agar conditions for 21 days. Crystal violet staining was performed, and the number of colonies were counted. The left panel shows a representative image of each treatment and the right panel the quantification and standard deviation. **B, C)** MTT assays were performed in MDA-MB-453 **(B)** and BT-549 **(C)** treated with 2 nM Paclitaxel (Px), 20 μM Enzalutamide (Enza), 5 μM AI-10-104 or all the combinations for 72 hrs. Results are expressed as the percentage of cell viability relative to control treatment (DMSO).

**Supplementary Table S1**. Statistical differences for all the treatment combinations of the experiments performed in MDA-MB-453 in Figure 5.

**Supplementary Table S2**. Statistical differences for all the treatment combinations of the experiments performed in SUM-159PT in Figure 5.

**Supplementary Table S3**. Statistical differences for all the treatment combinations of the experiments performed in MDA-MB-453 in Figure 6.

**Supplementary Table S4**. Primer’ sequences used for qPCR **(a)** or RUNX1 ChIP assay in MDA-MB-231 **(b)**.

