## Supplementary Figures and Tables for "RUNX1 is regulated by androgen receptor to promote cancer stem markers and chemotherapy resistance in triple negative breast cancer"

#### Supplementary Figure S1

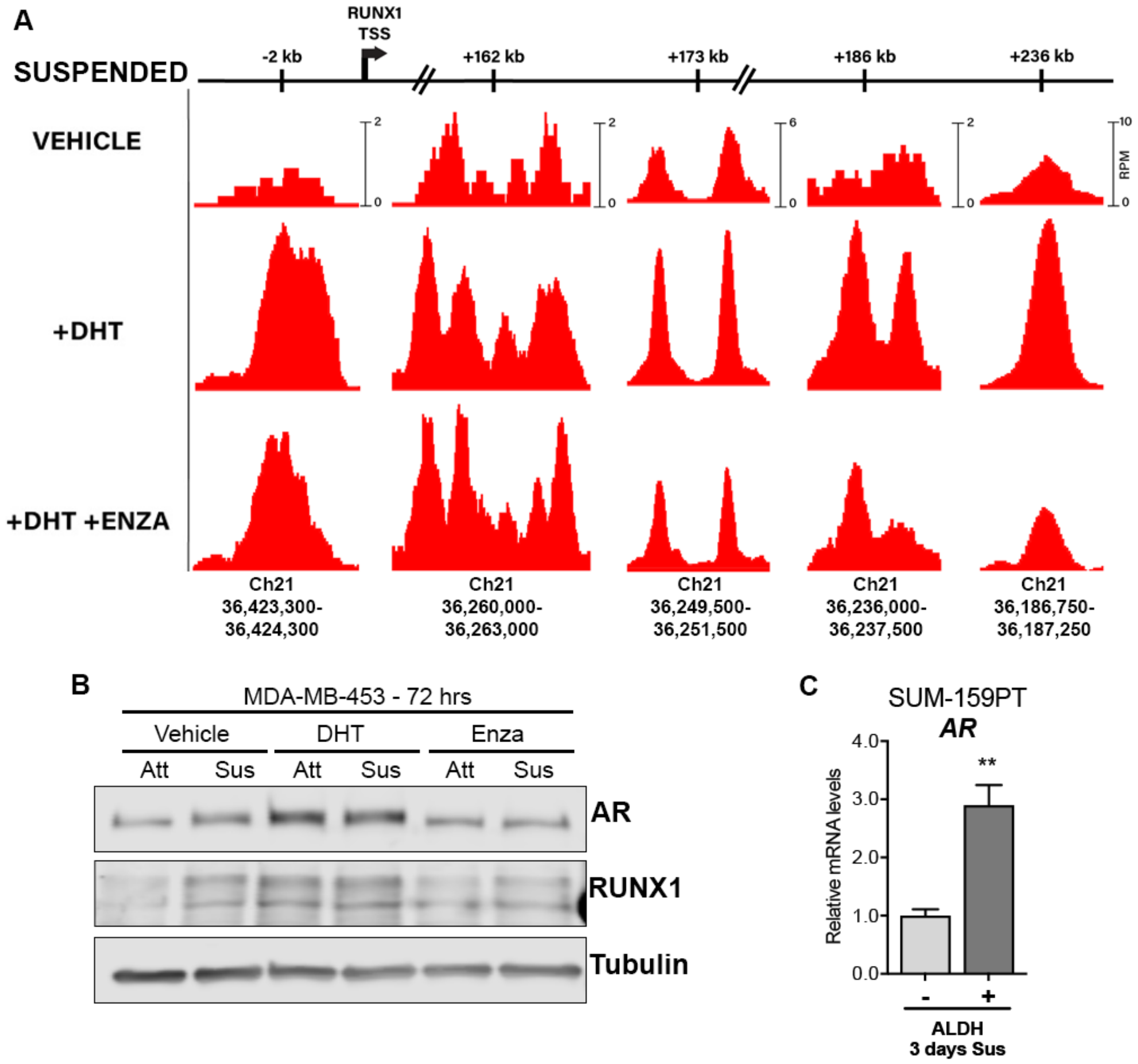

### Supplementary Figure S2

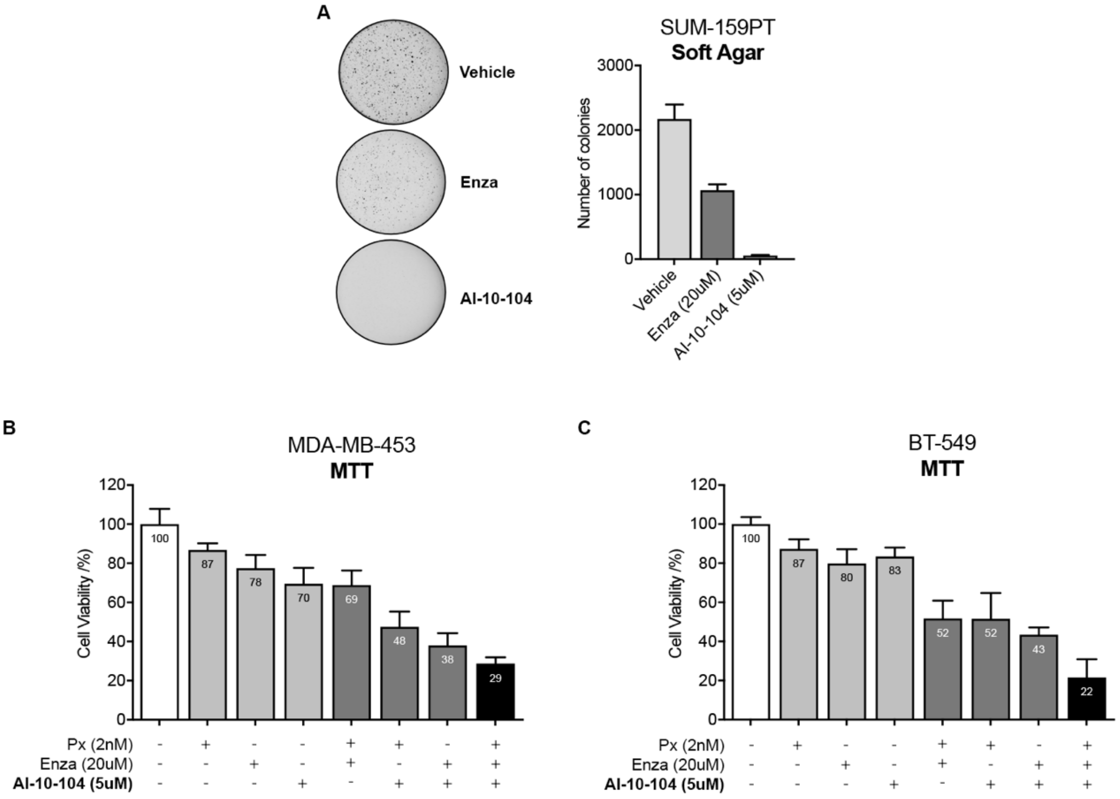

**Supplementary Table S1**. MDA-MB-453 Tukey's multiple comparisons test.

| Treatment comparison | Summary | Adjusted P Value |
|---|---|---|
| Vehicle vs. Px 1nM | ns | 0,1045 |
| Vehicle vs. Enza 20uM | ns | 0,1843 |
| Vehicle vs. AI-10-104 1uM | ** | 0,002 |
| Vehicle vs. Px + Enza | * | 0,0183 |
| Vehicle vs. Px + AI-10-104 | ** | 0,001 |
| Vehicle vs. Enza + AI-10-104 | ** | 0,0022 |
| Vehicle vs. Px + Enza + AI-10-104 | ** | 0,0011 |
| Px 1nM vs. Px + Enza | ns | 0,0863 |
| Px 1nM vs. Px + AI-10-104 | * | 0,0106 |
| Px 1nM vs. Px + Enza + AI-10-104 | ** | 0,0031 |
| Enza 20uM vs. Px + Enza | ns | 0,1041 |
| Enza 20uM vs. Enza + AI-10-104 | ** | 0,0057 |
| Enza 20uM vs. Px + Enza + AI-10-104 | ** | 0,0021 |
| AI-10-104 1uM vs. Px + AI-10-104 | ns | 0,2157 |
| AI-10-104 1uM vs. Enza + AI-10-104 | * | 0,0144 |
| AI-10-104 1uM vs. Px + Enza + AI-10-104 | ** | 0,0076 |
| Px + Enza vs. Px + Enza + AI-10-104 | * | 0,0267 |
| Px + AI-10-104 vs. Px + Enza + AI-10-104 | * | 0,0146 |
| Enza + AI-10-104 vs. Px + Enza + AI-10-104 | * | 0,0269 |

| Treatment comparison | Summary | Adjusted P Value |
|---|---|---|
| Vehicle vs. Px 1nM | ns | 0,2159 |
| Vehicle vs. Enza 20uM | ns | 0,1599 |
| Vehicle vs. AI-10-49 0.25uM | ns | 0,1751 |
| Vehicle vs. Px + Enza | * | 0,0443 |
| Vehicle vs. Px + AI-10-49 | * | 0,022 |
| Vehicle vs. Enza + AI-10-49 | *** | 0,0001 |
| Vehicle vs. Px + Enza + AI-10-49 | *** | 0,0004 |
| Px 1nM vs. Px + Enza | * | 0,0339 |
| Px 1nM vs. Px + AI-10-49 | ns | 0,1561 |
| Px 1nM vs. Px + Enza + AI-10-49 | * | 0,0121 |
| Enza 20uM vs. Px + Enza | ns | 0,1417 |
| Enza 20uM vs. Enza + AI-10-49 | ns | 0,1381 |

| | | |
|---|---|---|
| Enza 20uM vs. Px + Enza + AI-10-49 | * | 0,0152 |
| AI-10-49 0.25uM vs. Px + AI-10-49 | ns | 0,0606 |
| AI-10-49 0.25uM vs. Enza + AI-10-49 | * | 0,0374 |
| AI-10-49 0.25uM vs. Px + Enza + AI-10-49 | ** | 0,0028 |
| Px + Enza vs. Px + Enza + AI-10-49 | * | 0,0286 |
| Px + AI-10-49 vs. Px + Enza + AI-10-49 | ** | 0,0035 |
| Enza + AI-10-49 vs. Px + Enza + AI-10-49 | * | 0,039 |

**Supplementary Table S2**. SUM-159PT Tukey's multiple comparisons test.

| Treatment comparison | Summary | Adjusted P Value |
|---|---|---|
| Vehicle vs. Px 1nM | ns | 0,0966 |
| Vehicle vs. Enza 20uM | ns | 0,3025 |
| Vehicle vs. AI-10-104 1uM | ** | 0,0028 |
| Vehicle vs. Px + Enza | * | 0,0185 |
| Vehicle vs. Px + AI-10-104 | * | 0,0215 |
| Vehicle vs. Enza + AI-10-104 | ** | 0,0076 |
| Vehicle vs. Px + Enza + AI-10-104 | **** | <0.0001 |
| Px 1nM vs. Px + Enza | ns | 0,1998 |
| Px 1nM vs. Px + AI-10-104 | * | 0,0453 |
| Px 1nM vs. Px + Enza + AI-10-104 | ** | 0,0084 |
| Enza 20uM vs. Px + Enza | ns | 0,1977 |
| Enza 20uM vs. Enza + AI-10-104 | * | 0,042 |
| Enza 20uM vs. Px + Enza + AI-10-104 | ** | 0,0069 |
| AI-10-104 1uM vs. Px + AI-10-104 | ns | 0,1396 |
| AI-10-104 1uM vs. Enza + AI-10-104 | * | 0,0456 |
| AI-10-104 1uM vs. Px + Enza + AI-10-104 | ** | 0,0086 |
| Px + Enza vs. Px + Enza + AI-10-104 | * | 0,0218 |
| Px + AI-10-104 vs. Px + Enza + AI-10-104 | ns | 0,2231 |
| Enza + AI-10-104 vs. Px + Enza + AI-10-104 | ns | 0,1847 |

| Treatment comparison | Summary | Adjusted P Value |
|---|---|---|
| Vehicle vs. Px 1nM | ns | 0,2655 |
| Vehicle vs. Enza 20uM | ns | 0,1368 |

| | | |
|---|---|---|
| Vehicle vs. AI-10-49 0.25uM | ns | 0,1932 |
| Vehicle vs. Px + Enza | * | 0,0158 |
| Vehicle vs. Px + AI-10-49 | * | 0,018 |
| Vehicle vs. Enza + AI-10-49 | * | 0,0499 |
| Vehicle vs. Px + Enza + AI-10-49 | ** | 0,006 |
| Px 1nM vs. Px + Enza | ns | 0,1181 |
| Px 1nM vs. Px + AI-10-49 | ns | 0,1223 |
| Px 1nM vs. Px + Enza + AI-10-49 | * | 0,0253 |
| Enza 20uM vs. Px + Enza | ns | 0,2323 |
| Enza 20uM vs. Enza + AI-10-49 | ns | 0,1183 |
| Enza 20uM vs. Px + Enza + AI-10-49 | * | 0,012 |
| AI-10-49 0.25uM vs. Px + AI-10-49 | ns | 0,1693 |
| AI-10-49 0.25uM vs. Enza + AI-10-49 | ns | 0,4353 |
| AI-10-49 0.25uM vs. Px + Enza + AI-10-49 | ns | 0,0767 |
| Px + Enza vs. Px + Enza + AI-10-49 | ns | 0,171 |
| Px + AI-10-49 vs. Px + Enza + AI-10-49 | ns | 0,2302 |
| Enza + AI-10-49 vs. Px + Enza + AI-10-49 | ns | 0,0547 |

**Supplementary Table S3.** MDA-MB-453 Tukey's multiple comparisons test.

| Treatment comparison | Summary | Adjusted P Value |
|---|---|---|
| Vehicle vs. Pac 10nM | ns | 0,2241 |
| Vehicle vs. Enza 40uM | ns | 0,9263 |
| Vehicle vs. AI-10-104 5uM | ns | 0,2241 |
| Vehicle vs. AI-10-49 1uM | ns | 0,148 |
| **Vehicle vs. Pac + Enza + 104** | **** | <0.0001 |
| **Vehicle vs. Pac + Enza + 49** | **** | <0.0001 |
| Vehicle vs. Pac + Enza + 104 Att | ns | 0,5277 |
| Vehicle vs. Pac + Enza + 49 Att | ns | 0,577 |
| Pac 10nM vs. Enza 40uM | ns | 0,8843 |
| Pac 10nM vs. AI-10-104 5uM | ns | >0.9999 |
| Pac 10nM vs. AI-10-49 1uM | ns | >0.9999 |
| **Pac 10nM vs. Pac + Enza + 104** | ** | 0,0045 |
| **Pac 10nM vs. Pac + Enza + 49** | * | 0,0174 |

| | | |
|---|---|---|
| Pac 10nM vs. Pac + Enza + 104 Att | ns | 0,9992 |
| Pac 10nM vs. Pac + Enza + 49 Att | ns | 0,9982 |
| Enza 40uM vs. AI-10-104 5uM | ns | 0,8843 |
| Enza 40uM vs. AI-10-49 1uM | ns | 0,7693 |
| **Enza 40uM vs. Pac + Enza + 104** | *** | 0,0003 |
| **Enza 40uM vs. Pac + Enza + 49** | *** | 0,001 |
| Enza 40uM vs. Pac + Enza + 104 Att | ns | 0,9961 |
| Enza 40uM vs. Pac + Enza + 49 Att | ns | 0,9982 |
| AI-10-104 5uM vs. AI-10-49 1uM | ns | >0.9999 |
| **AI-10-104 5uM vs. Pac + Enza + 104** | ** | 0,0045 |
| AI-10-104 5uM vs. Pac + Enza + 104 Att | ns | 0,9992 |
| **AI-10-49 1uM vs. Pac + Enza + 49** | * | 0,0286 |
| AI-10-49 1uM vs. Pac + Enza + 49 Att | ns | 0,9865 |
| **Pac + Enza + 104 vs. Pac + Enza + 104 Att** | ** | 0,0012 |
| **Pac + Enza + 49 vs. Pac + Enza + 49 Att** | ** | 0,0041 |

**Supplementary Table S4a.** Primers' sequences used for qPCR

| Gene Name | Forward 5' - 3' | Reverse 5' - 3' |
|---|---|---|
| ***KLF4*** | CGAACCCACACAGGTGAGAA | TACGGTAGTGCCTGGTCAGTTC |
| ***GAPDH*** | GCACCACCAACTGCTTAGCA | GTCTTCTGGGTGGCAGTGATG |
| ***OCT4*** | CAGGCCCGAAAGAGAAAGC | CCACACTGGACCACATGGT |
| ***RUNX1*** | ACTCGGCTGAGCTGAGAAATG | GACTTGCGGTGGGTTTGTG |

**Supplementary Table S4b.** Primers' sequences used for the ChIP assay performed in MDA-MB-231

| Gene Name | Forward 5' - 3' | Reverse 5' - 3' |
|---|---|---|
| ***KCTD*** | GCTTCTGGGAAAGAACTGACATTC | GCCTTCTGTGCAAACATCTGAC |
| ***SOX4*** | TGCTTTTACCCACAAAAACAG | GGCCCAGGTTGTCAGACTTA |
